# Horizontal gene transfer rate is not the primary determinant of observed antibiotic resistance frequencies in *Streptococcus pneumoniae*

**DOI:** 10.1101/656751

**Authors:** Sonja Lehtinen, Claire Chewapreecha, John Lees, William P. Hanage, Marc Lipsitch, Nicholas J. Croucher, Stephen D. Bentley, Paul Turner, Christophe Fraser, Rafał J. Mostowy

## Abstract

The extent to which evolution is constrained by the rate at which horizontal gene transfer (HGT) allows DNA to move between genetic lineages is an open question, which we address in the context of antibiotic resistance in *Streptococcus pneumoniae*. We analyze microbiological, genomic and epidemiological data from the largest-to-date sequenced pneumococcal carriage study in 955 infants from a refugee camp on the Thailand-Myanmar border. Using a unified framework, we simultaneously test prior hypotheses on rates of HGT and a key evolutionary covariate (duration of carriage) as determinants of resistance frequencies. We conclude that in this setting, there is only weak evidence for the rate of HGT playing a role in the evolutionary dynamics of resistance. Instead, observed resistance frequencies are best explained as the outcome of selection acting on a pool of variants, irrespective of the rate at which resistance determinants move between genetic lineages.

---

Horizontal gene transfer (HGT), the non-vertical movement of genetic information between organisms, allows evolutionary innovation to spread onto new genetic backgrounds. However, the ability to predict the extent to which the frequency of such transfer impacts evolutionary outcomes (i.e., observed allele frequencies) remains a challenge in evolutionary biology. Beijerinck’s dictum “Everything is everywhere, but the environment selects”^1^ pithily captures the hypothesis that population sizes are sufficiently large that evolution is essentially the deterministic outcome of competition between all possible variants. A competing hypothesis is that, in practice, competition is limited to the pool of available variants and this pool is constrained by the rate at which genetic innovation arises. For this second hypothesis, lineages’ success therefore depends on the rate at which they acquire this innovation. In bacteria, the rate of horizontal movement of genetic material between lineages is a key determinant of this acquisition rate.

The evolution of antibiotic resistance is an important and illuminating area in which to address this question, because these two views of evolution lead to very different interpretations of observed patterns of resistance – in particular the heterogeneous distribution of resistance determinants among bacterial lineages. The ‘genetics as limiting factor’ hypothesis posits that different lineages have different resistance levels because they are able to acquire resistance determinants at different rates. This view implies that antibiotic resistance is beneficial and will eventually spread ubiquitously, but that lineages with higher rates of horizontal gene gain have been able to acquire these genes faster^2^. Conversely, the determinist evolutionary ecology perspective (‘everything is everywhere’) explains the distribution of resistance genes on lineages in terms of selection, that is, as the result of these genes conferring different fitness benefits on different lineages (e.g. Lehtinen et al.^3^ and San Millan^4^). Although lineages require HGT (or mutation) to acquire resistance determinants, it is assumed that these events occur often enough in the population so as not to limit the pool of available variants that selection can act on. Previously, these potential predictors of resistance have been considered separately; there is therefore need to revisit these hypotheses in a unified analysis. Here, we do so in the largest available population survey of *Streptococcus pneumoniae* that includes microbiological, genetic and epidemiological information about bacterial carriage.

In *S. pneumoniae*, both views are supported by large-scale data analyses^2,3,5^. A previously proposed determinist evolutionary ecology perspective of resistance evolution in *S. pneumoniae* explains variation in antibiotic resistance among pneumococcal lineages through variation in duration of carriage^3^, which itself is a heritable bacterial trait^6,7^. Theory suggests that the effect resistance has on the fitness of a lineage in species that are mainly carried asymptomatically, such as *S. pneumoniae*, depends on the lineage’s duration of carriage^3^. This is because these bacteria are primarily exposed to antibiotics prescribed for other infections (‘bystander selection’^8^) and the per-carriage-episode probability of antibiotic exposure is therefore greater for longer carriage episodes. Strains with a long duration of carriage thus gain more from being resistant to antibiotics. This model therefore predicts a positive association between resistance and duration of carriage in bacterial species like *S. pneumoniae*. The carriage duration of pneumococcal serotypes is indeed positively correlated with their resistance frequencies in multiple datasets^3^. The ‘genetics as limiting factor’ hypothesis, on the other hand, is supported by the observed positive association between levels of antibiotic resistance and recombination in the pneumococcus^2,9,10,11^.

The interpretation of these results is complicated by the observation that there is also an association between duration of carriage and HGT rate. Chaguza et al. find a positive correlation between the duration of carriage and HGT rate of pneumococcal serotypes^12^. This association raises the possibility that one of the other two associations could be confounded (Figure 1). In other words, it is possible that the association between resistance and HGT rate is not causal and arises because long duration of carriage causes both high resistance and high HGT rate. Similarly, it is also possible that the association between resistance and duration of carriage is itself not causal and arises because long duration of carriage causes high HGT rate, which in turn causes high resistance frequency.

**Figure 1:**
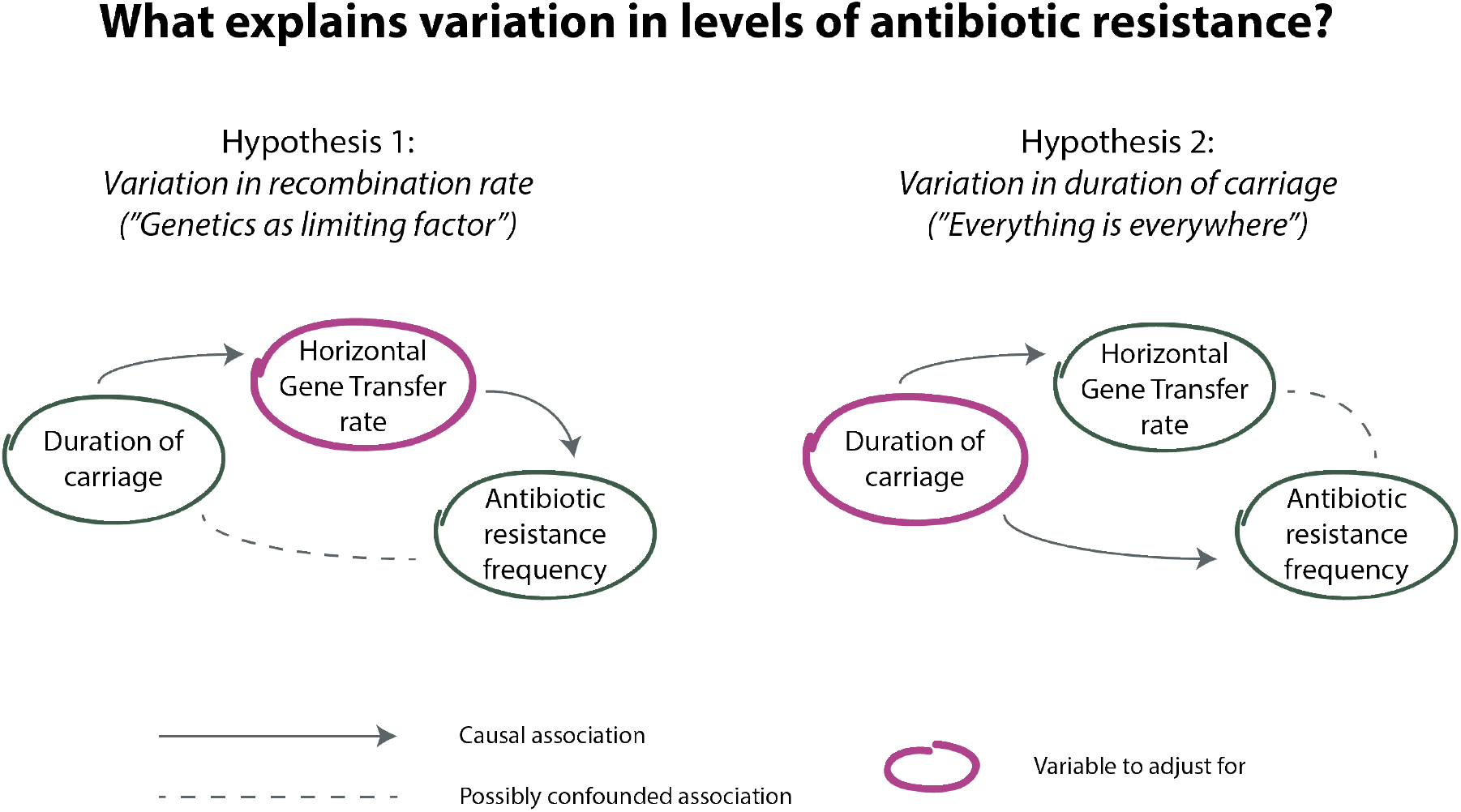
Schematic of the two possible explanations for variation in levels of antibiotic resistance between bacterial lineages. The first hypothesis is that lineages with high HGT rate have high resistance frequencies because they acquire resistance determinants at a higher rate. The second hypothesis is that lineages with a long duration of carriage have high resistance frequencies because resistance is more beneficial to these lineages (because longer duration of carriage translates into a greater probability of antibiotic exposure per carriage episode). Positive associations have been observed between all three variables (resistance, HGT rate and duration of carriage). If the first hypothesis is correct, the association between resistance and duration of carriage could be confounded by the causal path through HGT rate. If the second hypothesis is correct, the association between resistance and HGT rate could be confounded by the causal path through duration of carriage; in this case, these associations will not be robust to adjusting for the confounded variable.

In this paper, using a large pneumococcal isolate collection which combines epidemiological and genetic data, we investigated the relationship between antibiotic resistance, HGT rate and duration of carriage in a unified analytic framework. By testing which of the two properties of interest – HGT rate or duration of carriage – is the better predictor of per-lineage antibiotic resistance frequency, we assessed the relative importance of the ‘everything is everywhere’ and ‘genetics as limiting factor’ perspectives in pneumococcal resistance dynamics. The data were collected as part of a longitudinal study of pneumococcal carriage in a refugee camp on the Thailand-Myanmar border between 2007 and 2010^13^. Duration estimates were obtained for 1331 carriage episodes from Lees et al.^7^, while phenotypic data on resistance to seven different antibiotics were obtained from both Chewapreecha et al. and Lees et al.^14,7^. Whole-genome sequence data were obtained from 2663 isolates and classified into 48 groups of genetically related sequence clusters (SC), as described in Corander et al.^5^. HGT rate was quantified using two metrics: homologous recombination (HR), defined as the ratio of horizontally to vertically acquired single nucleotide polymorphisms (commonly known as *r/m*); and gene movement (GM) defined as the sum of gene gains or losses per vertical substitution. These two measure aim to capture different mechanisms of HGT: in *S. pneumoniae* HR quantifies transformation, the incorporation of fragments of DNA through homologous recombination, whereas GM captures the movement of integrative conjugative elements. The two metrics are not correlated (Pearson’s correlation coefficient: −0.09, 95% CI [−0.37,0.20]), suggesting they indeed capture different facets of HGT (see Supporting Information). We used these data to test the association between i) resistance and duration of carriage and ii) resistance and HGT rate, controlling for confounding with the other variable.

Testing for association at the level of individual carriage episodes is problematic because those carriage episodes may be closely related via recent transmission and may therefore be non-independent. To circumvent this problem, episodes of carriage need to be aggregated prior to testing for association. Episodes of carriage could be aggregated by serotype or sequence cluster (SC). The choice of aggregator is not trivial: both serotype and SC are potentially predictors of our three traits of interest. We therefore used regression models to test which aggregator is the best predictor of the three traits of interest and found that models with SC as the predictor fit the data best (see Table 1 and Supporting Information).

**Table 1:** Akaike Information Criterion for regression models using serotype, SC and serotype-SC combination as a predictor of resistance, HGT rate and duration of carriage (see Supporting Information). The lowest AIC is indicated in bold. SC = sequence cluster, HR = homologous recombination, GM = gene movement.

| Trait | serotype | SC | serotype-SC |
| --- | --- | --- | --- |
| Resistance multiplicity | 3224 | **3163** | 3191 |
| Carriage duration | 13243 | **13220** | 13269 |
| HR | 4370 | 4271 | **4251** |
| GM | **88928** | 88960 | 88965 |

We therefore averaged the data by SC, and computed associations between resistance (both in terms of resistance multiplicity, the number of antibiotics an isolate is resistant to, and individually against each antibiotic) and HGT rate and duration of carriage using Kendall’s rank correlation coefficient *τ*. Confidence intervals (CIs) were generated using bootstrapping (see Supporting Information). We found strong evidence for an association between resistance multiplicity and duration of carriage (*τ* : 0.41, 95%CI [0.16,0.60]), which is robust to adjusting for either measure of HGT rate (Figure 2). We found no evidence for an association between GM and resistance multiplicity, whether or not adjusting for duration of carriage. We found weak evidence for an association between resistance multiplicity and HR (*τ* : 0.20, 95%CI [0.00,0.38]), that became non-significant when adjusting for duration of carriage.

**Figure 2:**
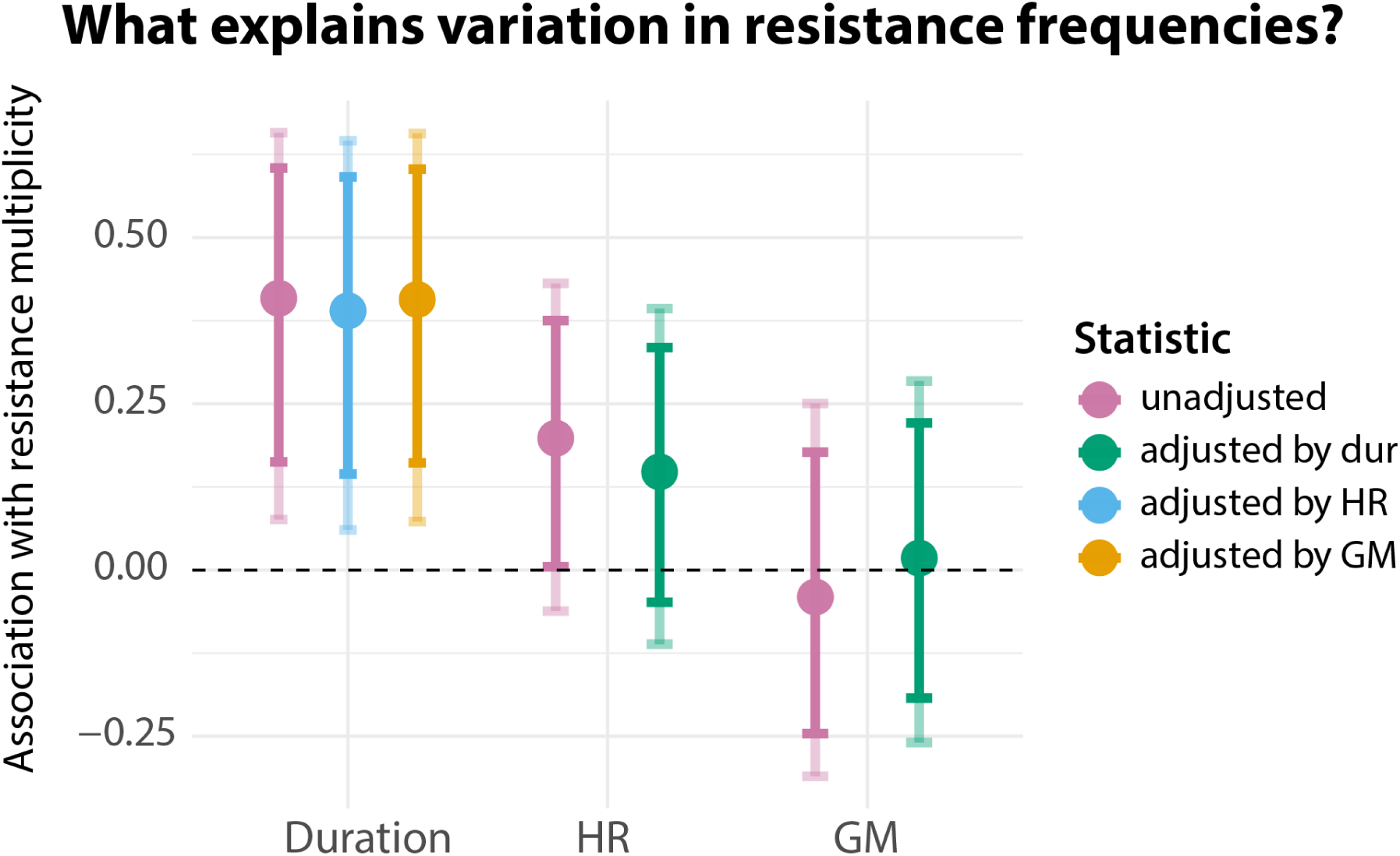
Kendall rank correlation (*τ*) between sequence cluster (SC) resistance multiplicity and duration of carriage (unadjusted and adjusted for HR or GM); SC resistance multiplicity and HR (unadjusted and adjusted for duration of carriage); and SC resistance multiplicity and GM (unadjusted and adjusted for duration of carriage). Error bars represent 95% confidence intervals and were computed by bootstrapping (see Supporting Information). SCs with fewer than 5 episodes of carriage were excluded, giving a sample size of 43 SCs.

Similar trends held when analysing antibiotics individually (Figure 3): in general, resistance was more strongly associated with duration of carriage than either measure of HGT rate and this association was undiminished when adjusting for HGT. Cotrimoxazole was an exception to this pattern: cotrimoxazole resistance was more strongly associated with HR than duration of carriage, and this relationship remained statistically significant when adjusting for duration of carriage. This is consistent with previous observations that cotrimoxazole resistance emerges via HR-mediated acquisition of alleles^15^ and the corresponding genes were found to be recombination hotspots^14^.

**Figure 3:**
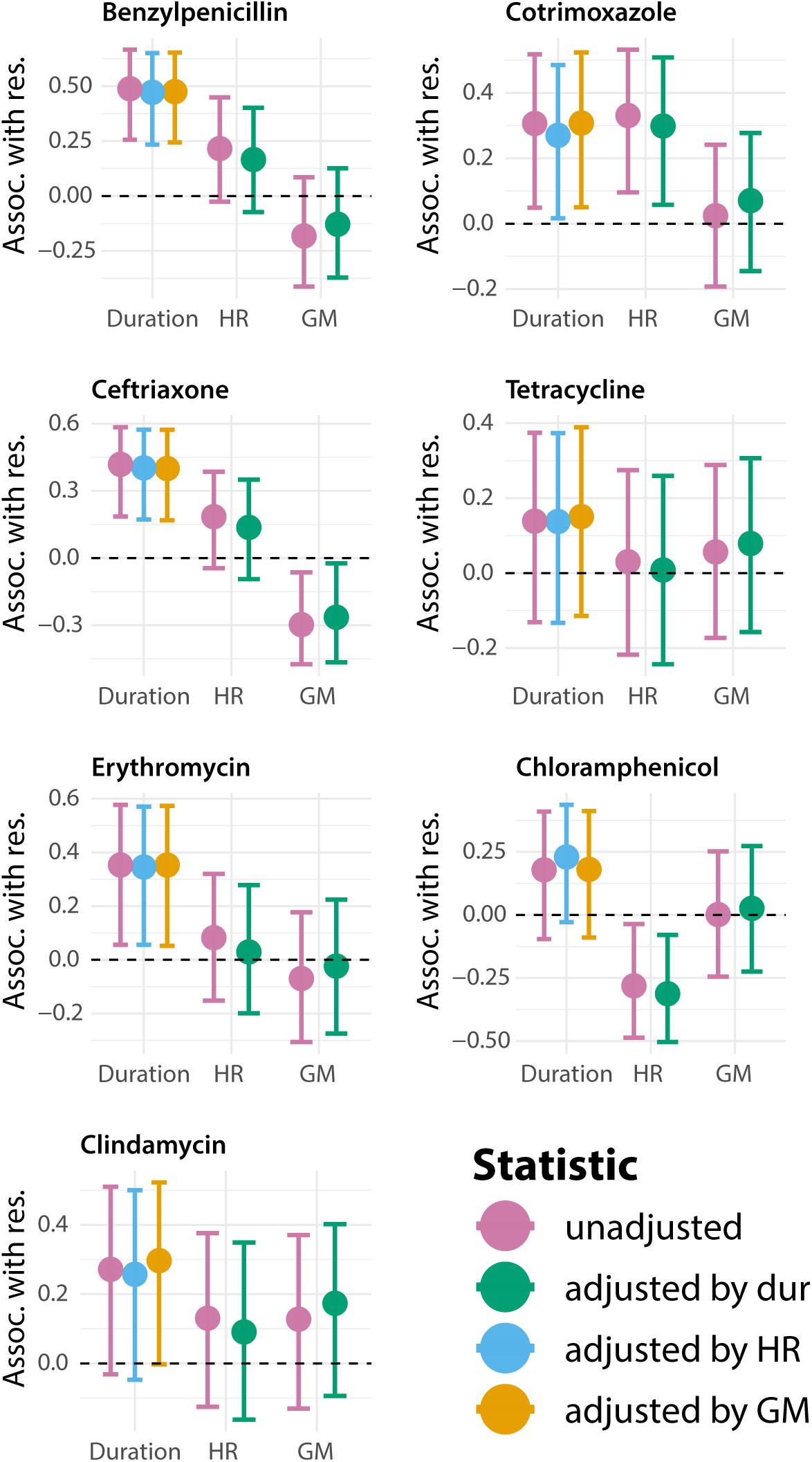
Kendall rank correlation (*τ*) between sequence cluster (SC) resistance to each antibiotic and duration of carriage (unadjusted and adjusted for HR or GM); SC resistance to each antibiotic and HR (unadjusted and adjusted for duration of carriage); and SC resistance to each antibiotic and GM (unadjusted and adjusted for duration of carriage). Opaque error bars represent 95% confidence intervals while transparent error bars represent 99% confidence intervals; both were computed by bootstrapping (see Supporting Information). SCs with fewer than 5 episodes of carriage were excluded, giving a sample size of 42 SCs.

Results of the analysis were qualitatively similar when using serotype or serotype-SC as the analysis aggregators (see Figures S2 and S3). First, the association between resistance multiplicity and carriage duration remained significant, even when adjusting for either measure of HGT. Second, the association between resistance and GM remained non-significant with and without adjusting for the duration of carriage. We did not find strong evidence for the association between resistance and HR when adjusting for the duration of carriage: this association was significant when using serotype-SC as the aggregator (*τ* : 0.18, 95%CI [0.01,0.33]), and was non-significant when using serotype as the aggregator.

Finally, we did not replicate a significant correlation between HGT rate and duration of carriage for either of our measures of HGT rate (*τ* : 0.15, 95%CI [−0.11,0.40] for HR; −0.14, 95%CI [–0.37,0.11] for GM). These associations remained non-significant in the case of the two remaining aggregators, serotype and serotype-SC.

Taken together, these results provide strong support for an association between duration of carriage and resistance, which is robust to adjusting for the rate of HGT. Conversely, we found only weak evidence for an association of homologous recombination and resistance, which weakened further when adjusting for duration of carriage; we found no evidence for an association between gene movement and resistance. Our data thus do not support a major role for HGT rate in determining the frequency of antibiotic resistance in pneumococcal lineages, hence lending support to the deterministic eco-evolutionary perspective.

This work was motivated by the reported correlations between resistance, HGT rate (specifically HR) and duration of carriage. We only replicate a significant correlation for resistance and duration of carriage – our point estimates for the other two correlations are positive, but generally non-significant. One possible explanation for this discrepancy could be differences in the way HR is estimated. This is unlikely to be the case, however, as our estimates are strongly correlated with previously used measures (Supplementary Figure S1). Another potential explanation is that our treatment of uncertainty is more conservative than in previous work. Our confidence intervals take into account the uncertainty in estimating the per-cluster averages. This is particularly significant for HR, because recombination rates are often based on a small number of recombination events, leading to large confidence intervals and thus potentially explaining why we observe positive, but non-significant, associations between HR and resistance, and HR and duration of carriage (Supplementary Figure S2). Indeed, computing the correlation between HR and duration of carriage without accounting for uncertainty in the per-cluster estimates gives a higher, though still negative, lower bound for the 95% confidence interval (Supplementary Table S2).

Point estimates for the association between HR and resistance were consistently positive (though non-significant), even when adjusting for duration of carriage. Given our conservative treatment of uncertainty in the estimates of HR, these results do not rule out a potential independent role for HR, or some other correlate of HR, in determining resistance frequencies. Indeed, a potential causal role for HR is supported by the observation that the magnitude of the correlation between HR and resistance was greatest for the resistance determinants that are not carried on transposons and would therefore be expected to be acquired through HR rather than GM (benzylpenicillin, ceftriaxone, cotrimoxazole resistance). Nevertheless, although HR rate cannot be ruled out as a driver of resistance frequencies, the small magnitude of the association between HR and resistance suggests that this role is relatively minor compared to the role of duration of carriage.

The robustness of the association between duration of carriage and resistance supports the role of bacterial duration of carriage as a major driver of antibiotic resistance. An important caveat here is that there is uncertainty with respect to the direction of causality between resistance and duration of carriage; such association could also arise because resistance prevents clearance through antibiotic exposure thus prolonging the duration of carriage. We attempted to establish whether the association between resistance and carriage duration was still present when eliminating the effect of resistance on duration of carriage, but our results were inconclusive (presumably due to insufficient statistical power, see Supplementary Text). However, as some of us have previously suggested^16^, the association being observed at the serotype level lends support to the role of duration of carriage in determining resistance frequencies.

In conclusion, our analysis supports an ‘everything is everywhere’ view of resistance evolution, in which observed allele frequencies and combinations depend on the fitness of these genotypes and are not constrained by a history of genetic events. Yet, the role of HGT in acquisition of resistance determinants is well documented^17,18,14,19,15^. This apparent contrast can be resolved by placing our results in the context of the time frame of this study. Antibiotic use has been present over a timescale of several decades, and, during the course of this study, resistance frequencies were not observed to change (see Supplementary Figure S7), suggesting they were probably at quasi-equilibrium. The picture that emerges, therefore, is that while HGT rate plays a role in how long it takes for a pneumococcal populations to acquire novel resistance determinants, the eventual frequency of these determinants can be understood purely in terms of evolutionary optimisation. Our study leaves open the question of the role of genetic processes in non-equilibrium populations. Public health measures such as vaccination programs or changes in treatment regimes can be viewed as perturbations that change the fitness landscape of the pathogen population. Addressing the relative importance of genetic and selection processes in the short term dynamics following such a perturbation is therefore necessary to predict the effect of medical interventions on bacterial populations.

## Acknowledgements

This work was funded by the Imperial College Research Fellowship (RM), the Polish National Agency of Academic Exchange (RM), the National Institutes of Health (MIDAS) (SL and CF grant ref: U01GM110721-01), the Li Ka Shing foundation (SL and CF), the Sir Henry Wellcome postdoctoral Fellowship (CC, grant ref: 107376/Z/15/Z), the National Institutes of Health (WPH, grant ref: NIH R01 AI106786), a Sir Henry Dale Fellowship, jointly funded by Wellcome and the Royal Society (NJC, grant ref: 104169/Z/14/Z), and the National Institute Of General Medical Sciences (ML, cooperative agreement U54GM088558). The content is solely the responsibility of the authors and does not necessarily represent the official views of the National Institute Of General Medical Sciences or the National Institutes of Health.

