## Supplementary material for "Horizontal gene transfer rate is not the primary determinant of observed antibiotic resistance frequencies in *Streptococcus pneumoniae*": supporting-information.pdf

#### 1 Supplementary Text

##### 1.1 Episodes of carriage: duration, resistance and cluster assignment

###### 1.1.1 Data sources

Data used in this analysis were collected as part of a longitudinal study of pneumococcal carriage in infants and their mothers in a 4 $km^2$-area refugee camp on the Thailand-Myanmar border area between 2007 and 2010, where the pneumococcal vaccine was not available [1, 2]. Briefly, the study recruited 999 pregnant women and followed all 955 live births until their second birthday. Nasopharyngeal swabs were taken monthly from all infants and from a subset of 234 mothers. All swabs from the 234 mother-infant pairs, as well as selected swabs from the remaining infants (those taken during or immediately prior to a clinical pneumonia episode), were cultured and serotyped according to the World Health Organisation protocol as described in [1] and phenotypically tested for susceptibility to seven classes of antibiotics (benzylpenicillin, ceftriaxone, cotrimoxazole, tetracycline, erythromycin, chloramphenicol and clindamycin) and classified as susceptible, intermediate or resistant according to the Clinical and Laboratory Standards Institute criteria (2007). Intermediate isolates were considered resistant. A random subset of these samples were subsequently whole-genome sequenced, as described in [3]. These data ($n = 2663$ genome assemblies) were obtained from Corander et al. [4]. Additionally, all swabs from another 364 infants (i.e. infants not part of the 234 mother-infant pairs) were serotyped using the latex sweep method as described in [5]. The two longitudinal datasets (i.e. swabs from the 234 infants and 364 infants) had previously been used to estimate duration of different episodes of carriage by fitting a hidden Markov model [6]. These carriage estimates ($n = 1331$ episodes) were obtained from Lees et al [6].

###### 1.1.2 Serotype and cluster assignment

We used the pipeline previously described in Mostowy et al. [7] to infer serotype based on genetic sequence. Sequence quality was too low to infer serotype for 4 of the isolates; these isolates were assigned the experimentally characterised serotype. For a small number of swabs (77), the experimentally determined serotype was different from the sequence-derived serotype; for these we used the sequence-derived serotype for further analysis. Sequence cluster (SC) assignments were obtained from Corander et al. [4]. These had been determined using a species-wide clustering analysis based on genetic similarity combining four major datasets. In order to obtain reliable estimates of horizontal gene transfer rate (see below), SCs with below 10 isolates were excluded from further analysis, leaving 48 out of 55 SCs.

###### 1.1.3 Combining datasets

Combining the genetic and epidemiological datasets gave a set of carriage episodes ($n = 1179$) with associated duration, serotype, resistance to seven classes of antibiotics and SC assignment. When multiple sequences originated from the same carriage episode, these were considered as a single data point. On occasion, sequences from the same carriage episode had inconsistent serotypes (7 carriage episodes) or SC assignments (14 carriage episodes). Inconsistency in serotype was due to inconsistency between genetically and experimentally inferred serotype. Inconsistency in SC assignment was due to episodes of carriage having been inferred based on serotype only: two consecutive episodes of carriage of genetically distinct strains with the same serotype would have been inferred to be a single episode of carriage. These episodes of carriage were removed, along with carriage episodes from clusters with fewer than 10 isolates (see above), giving a dataset of 1091 carriage episodes. For 145 of these, sequences from the same episode of carriage had consistent SC assignments but different resistance profiles. We assumed this represented within-host evolution rather than infection with distinct strains. For these episodes of carriage we used the mean value of the resistance statuses (e.g., if the episode consisted of one resistant and one susceptible isolate, we used 0.5 as the resistance status).

#### 1.2 Horizontal gene transfer rate

Horizontal gene transfer (HGT), broadly defined as horizontal transfer of genetic information, was quantified, independently for each SC, using two metrics. The first metric, called homologous recombination (HR), was the proportion genetic variation (single nucleotide polymorphisms – SNPs) acquired horizontally to genetic variation acquired vertically, commonly known as $r/m$. The second metric, called gene movement (GM), captures the rate of gene gain or loss in the SC. To obtain these measures, we predicted protein coding sequences in 2,663 genome assemblies using Prodigal [8], and assigned these sequences into clusters of orthologous genes (COGs) using `mmseqs` with default parameters and `--min-seq-id 0.5` option [9]. We next defined a core genome for each SC as a set of COGs in a given SC that are uniquely present in at least 70% of all isolates in that SC. These COGs were then concatenated into a core genome alignment and used as an input for Gubbins with default parameters, which uses a phylogenetic approach to assign SNPs into those acquired horizontally and vertically and returns a clonal phylogeny for each SC [10]. Next, we inferred the number of gains and losses of the remaining COGs on each branch of that clonal tree using GLOOME, which uses a stochastic mapping approach to estimate the horizontal gene flow given a gene presence/absence profile and a phylogenetic tree. The end result of this analysis was a table where each row showed an independent branch of a clonal tree in a given SC, the number of SNPs acquired via mutation and recombination, number of gene gains and gene losses, and the branch length following a removal of horizontally acquired variation. The longest branches of the tree were excluded due to large uncertainty in recombination estimates at those branches (5000 SNPs or longer). The HR rate was defined as the total sum of horizontally acquired SNPs to the sum of vertically acquired SNPs. The GM rate was defined as the total sum of gene gains and losses divided by the total branch length of the clonal tree. The confidence intervals were obtained by resampling branches of the tree with replacement $n = 1000$ times for each SC.

To validate our pangenome-based method of inferring HR rates, we applied this method to $n = 616$ isolates from a Massachusetts pneumococcal collection for which $r/m$ rates were obtained by mapping each SC to a draft pneumococcal genome (see Figure S1). Results show that, while there is a strong concordance between the two measures ($R^2 = 0.89$, 95% CI: [0.44,0.98]), there is a small discrepancy between the two measures for some sequence clusters. To investigate whether this discrepancy is due to a false detection rate of HGT stemming from sub-optimal concatenation of COGs in the core alignment, we investigated the effect of random COG concatenation on the estimation of HR and GM rates (see Figure S2). We found that the uncertainty in estimates of both HR and GM rate is comparable or smaller than the rate uncertainty obtained by branch resampling, suggesting that the COG concatenation has a limited influence on the estimate of HGT rates. We thus conclude that the observed discrepancies between the estimates of $r/m$ using our pangenome approach and those approaches using a full reference sequence are mostly driven by the gene movement of low-to-intermediate frequency genes in the reference sequence.

Finally, to confirm that our estimates of HGT rate were not driven by transfer events involving resistance genes (in which case, high HGT rates in a particular lineage might simply have reflected higher selection pressure for resistance), we re-estimated HR and GM rates having removed antibiotic resistance determinants (Figure S8). These estimates were strongly correlated with the estimates in the main analysis (XX and XX), suggesting that observed HGT rates are not driven by the acquistion of resistance.

#### 1.3 Choice of aggregator

The choice of aggregator was based on fitting regression models to determine whether serotype, SC or serotype-SC best predicted the traits of interest (HGT rate, duration of carriage and resistance status). For resistance multiplicity and duration of carriage, which are properties of a carriage episode, we fitted Poisson and linear regression models (respectively) to the carriage episode dataset. For the HGT rate metrics, the regression models (logistic for HR and linear for GM) were fitted to the phylogeny branches. The quality of the fits was assessed using AIC ($-2\ln(\hat{L})+2k$), where $\hat{L}$ is the maximum value of the model's likelihood function and $k$ is the number of parameters to fit (48, 48 and 86 for serotype, SC and serotype-SC respectively in the carriage episode dataset).

#### 1.4 Correlation with resistance

Associations between SC mean resistance and SC mean recombination rate, and SC mean resistance and SC mean duration of carriage, were tested using Kendall's rank correlation tau ($\tau$). Kendall's tau was chosen as a metric (rather than Spearman's rho) because of the nature of uncertainty in our data: clusters with fewer samples are likelier to have extreme resistance values (i.e. 0% or 100% resistance) and Kendall's tau is less sensitive to error at the extremes of the distribution than Spearman's rho. The adjusted associations between resistance and recombination rate controlling for carriage duration, and resistance and carriage duration controlling for recombination rate were calculated using partial Kendall's rank correlation as implemented in `ppcor` R package [11].

Calculating the correlation using SC means would not account for the uncertainty in the estimation of these means. To address this, we used a dual bootstrap approach to compute confidence intervals for tau: if $P(\tau|\theta)$ is the sampling distribution of tau, given the SC properties to correlate ($\theta$, e.g. resistance and duration of carriage) and $P(\theta)$ is the sampling distribution of the SC properties, the overall sampling distribution for $P(\tau)$ is $P(\tau) = \int_{\theta} P(\tau|\theta)P(\theta)d\theta$. Both $P(\tau|\theta)$ and $P(\theta)$ were estimated using bootstrapping using the following procedure:

1. For each SC, draw, with replacement, from carriage episodes in the SC, a set of carriage episodes of the same size as the SC. Calculate the mean carriage duration.

2. Similarly, for each SC, draw a set of resistance multiplicities and calculate mean resistance multiplicity.

3. For each SC, draw, with replacement, from the SC phylogeny, a set of random branches the same size as the number of branches in the SC phylogeny. Calculate mean HR and HGT.

4. Sample with replacement among these SC estimates, calculate $\tau$, repeat $m_b$ times to give estimate of $P(\tau|\theta)$ for a particular $\theta$.

5. Repeat steps 1-4, $n_b$ times.

6. Sum the $n$ distributions to give $P(\tau)$

95% confidence intervals were calculated from this distribution. SCs with fewer than five observations were excluded from the analysis. The bootstrap samples used are $n_b = m_b = 1000$ unless stated otherwise.

#### 1.5 Direction of causality for resistance and duration of carriage

As discussed in the main text, we expect a long duration of carriage to be predictive of resistance because resistance is more advantageous in lineages with a long duration of carriage. However, we also expect long duration of carriage to be associated with resistance because resistant strains are less likely to be cleared through antibiotic exposure. To test whether this reverse causality accounts for the association we observe, we compute per-SC carriage duration using only episodes of carriage with the same resistance profile. This eliminates the effect of between-SC variation in resistance frequency on duration of carriage. We use the most common resistance profile (resistance to cotrimoxazole and sensitivity to all other antibiotics, $n = 188$ carriage episodes), which reduces the number of SCs for which we can compute the duration of carriage, to $n = 11$. We find a positive, but no longer significant association between resistance and duration of carriage (**$\tau$: 0.25, 95%CI [-0.42,0.76]**).

The loss of significance might be due to the smaller number of SCs (as opposed to the elimination of the effect of resistance on duration of carriage). To check for this, we look at the correlation between resistance and duration of carriage for the same set of 11 SCs, this time using the duration of carriage estimates from the main analysis (i.e. without restricting to a single resistance profile). Again, we see a positive, but non-significant association **($\tau$: 0.41, 95%CI [-0.23,0.86])**. This suggests that the loss of the significance is at least partially due to the decrease in sample size. We therefore cannot confidently conclude whether the association between resistance and duration of carriage is driven by the effect of resistance on duration of carriage, the effect of duration of carriage on resistance, or a combination of the two.

### 2 Supplementary Tables

**Table S1.** Akaike Information Criterion for logistic regression model using serotype, cluster and serotype-SC combination as a predictor of resistance, HGT rate and duration of carriage (see Methods for details) for resistance against individual antibiotics. The lowest AIC is indicated in bold. The table can be found in a separate CSV file.

**Table S2.** SC averages for resistance multiplicity, individual resistance, duration of carriage, HR and GM, excluding SCs with fewer than 5 episodes of carriage. The table can be found in a separate CSV file.

### 3 Supplementary Figures

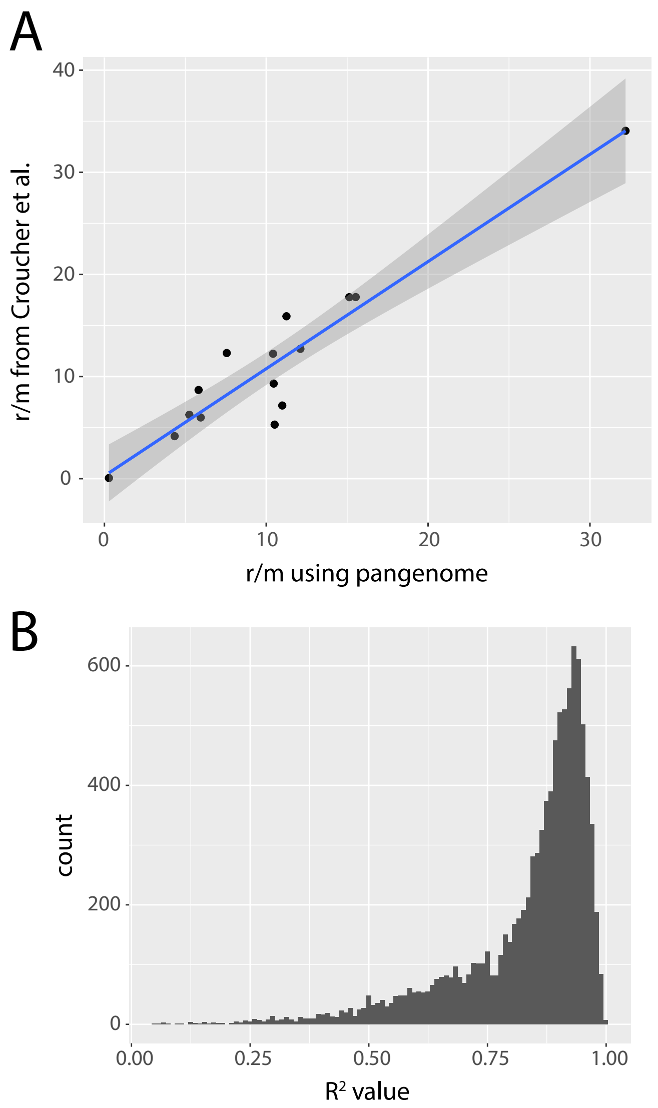


**Figure S1.** Validity of pangenome-based measure of homologous recombination. (A) X-axis shows the HR ($r/m$) rate obtained as described in the text for $n = 15$ SCs from Croucher et al. [12], while Y-axis shows the original estimates obtained by mapping short-reads to 15 corresponding reference draft genomes. (B) Histogram of $R^2$ values obtained by resampling 15 points in panel A with replacement $n = 1000$ times and reestimating the correlation coefficient. The resulting correlation between estimates from the two approaches is $R^2 = 0.89$, 95% CI: [0.44,0.98].

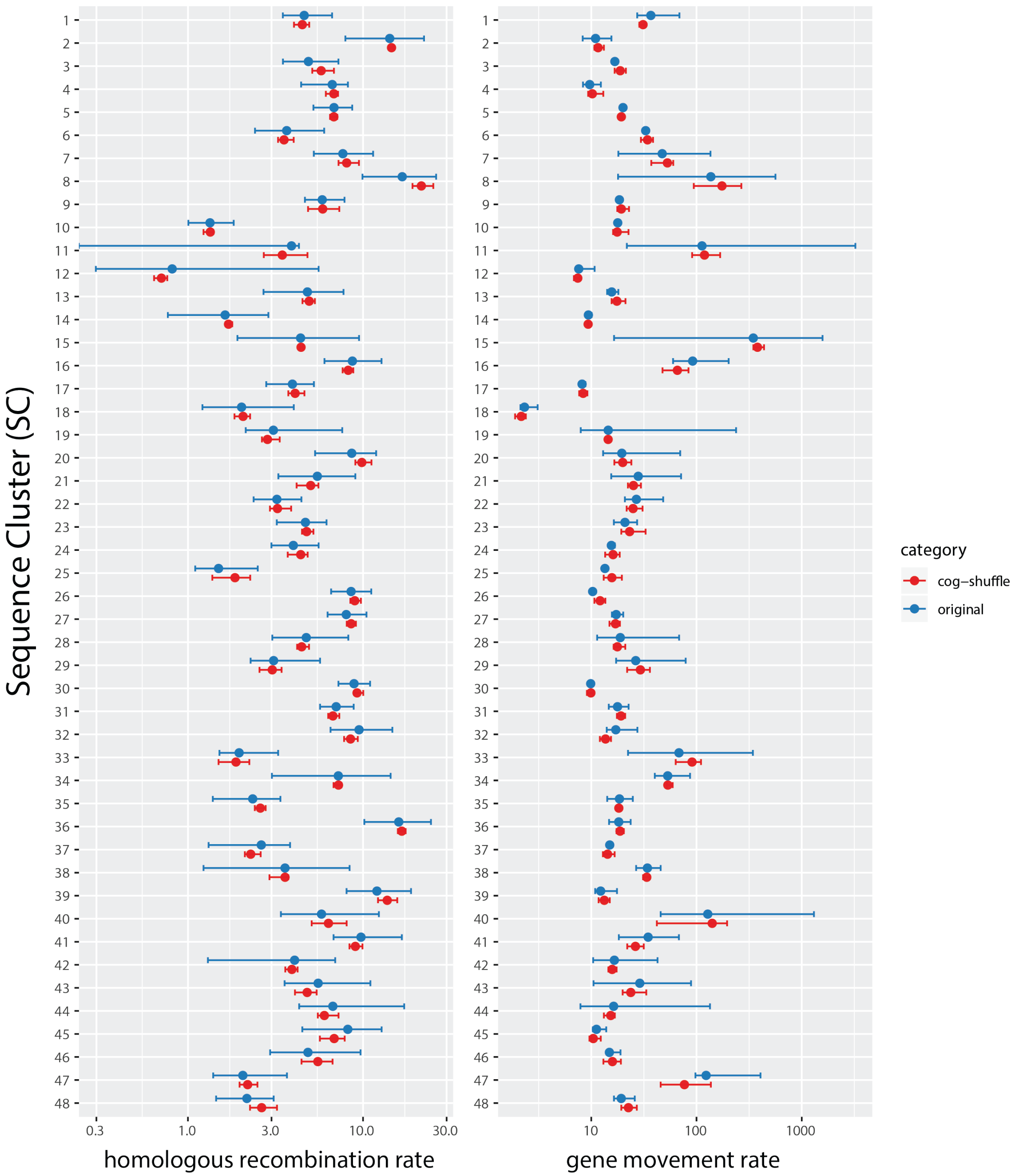


**Figure S2.** Effect of random order COG concatenation on the estimation of HR rate (left) and GM rate (right). Blue points show the main estimates for all 48 sequence clusters (SC), with error bars showing the 95% confidence intervals obtained by bootstraping branches of the clonal tree, as described in the Supplementary Text. Red error bards and points show the full range of HR/GM values and their median, respectively, obtained by shuffling COGs randomly.

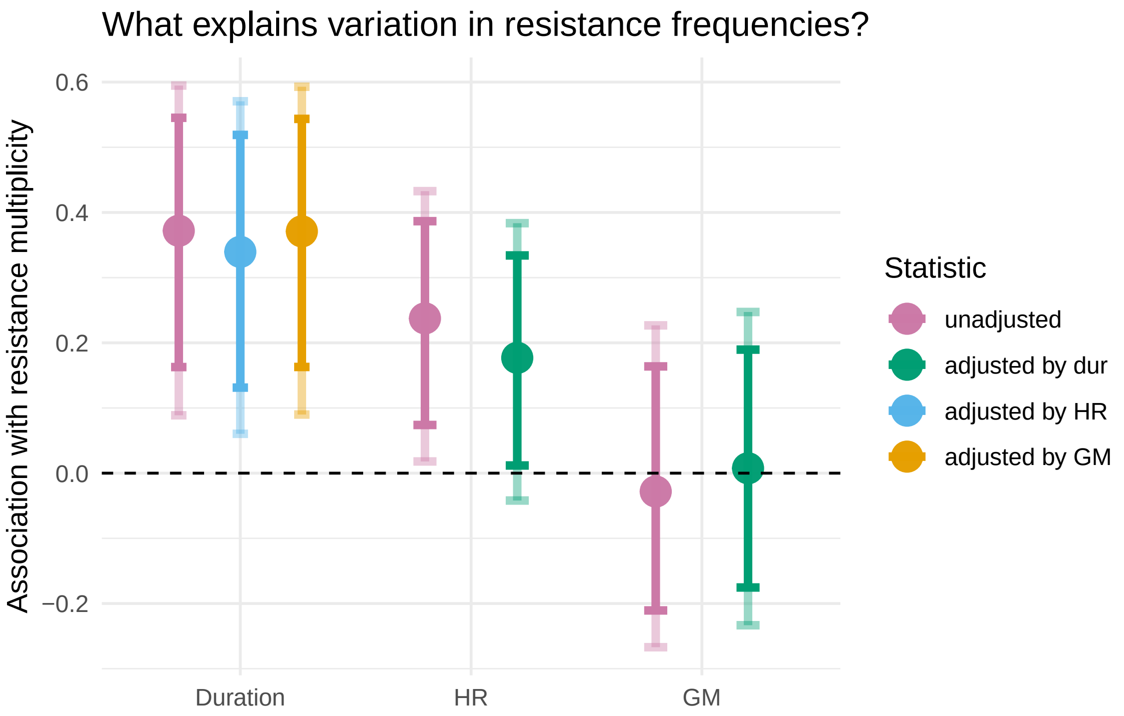


**Figure S3.** Association between resistance and duration of carriage and two measures of HGT, with serotype-SC as the main unit of analysis. The legend of each panel is the same as described in Figure 2.

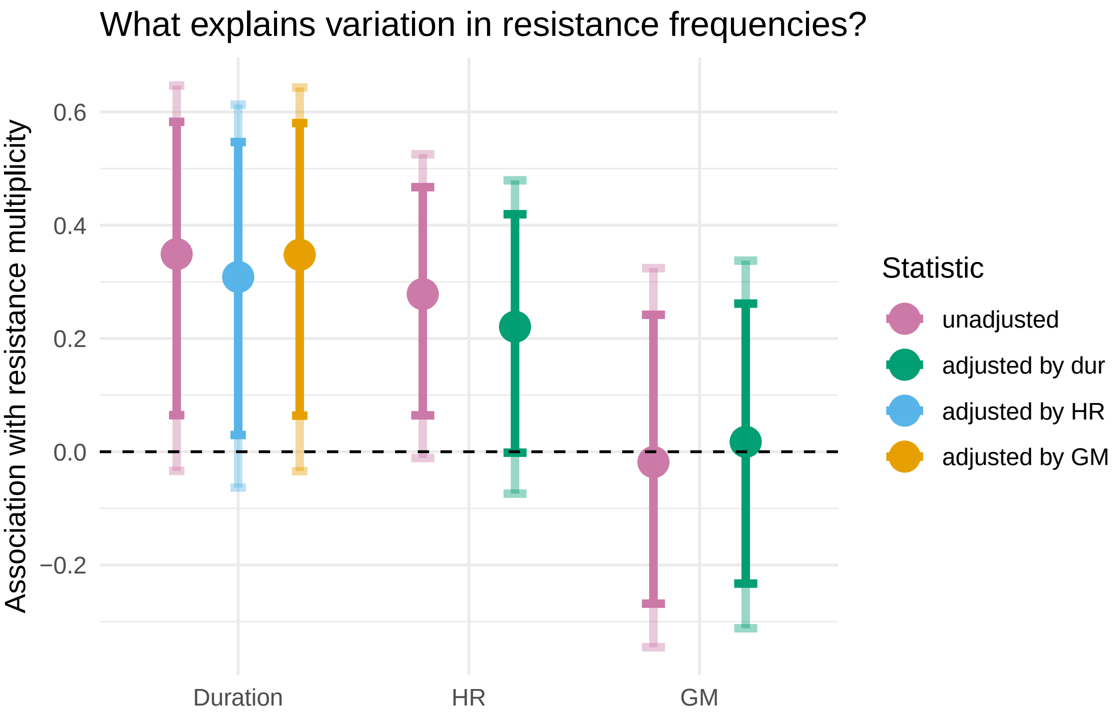


**Figure S4.** Association between resistance and duration of carriage and two measures of recombination, with serotype as the main unit of analysis. The legend of each panel is the same as described in Figure 2.

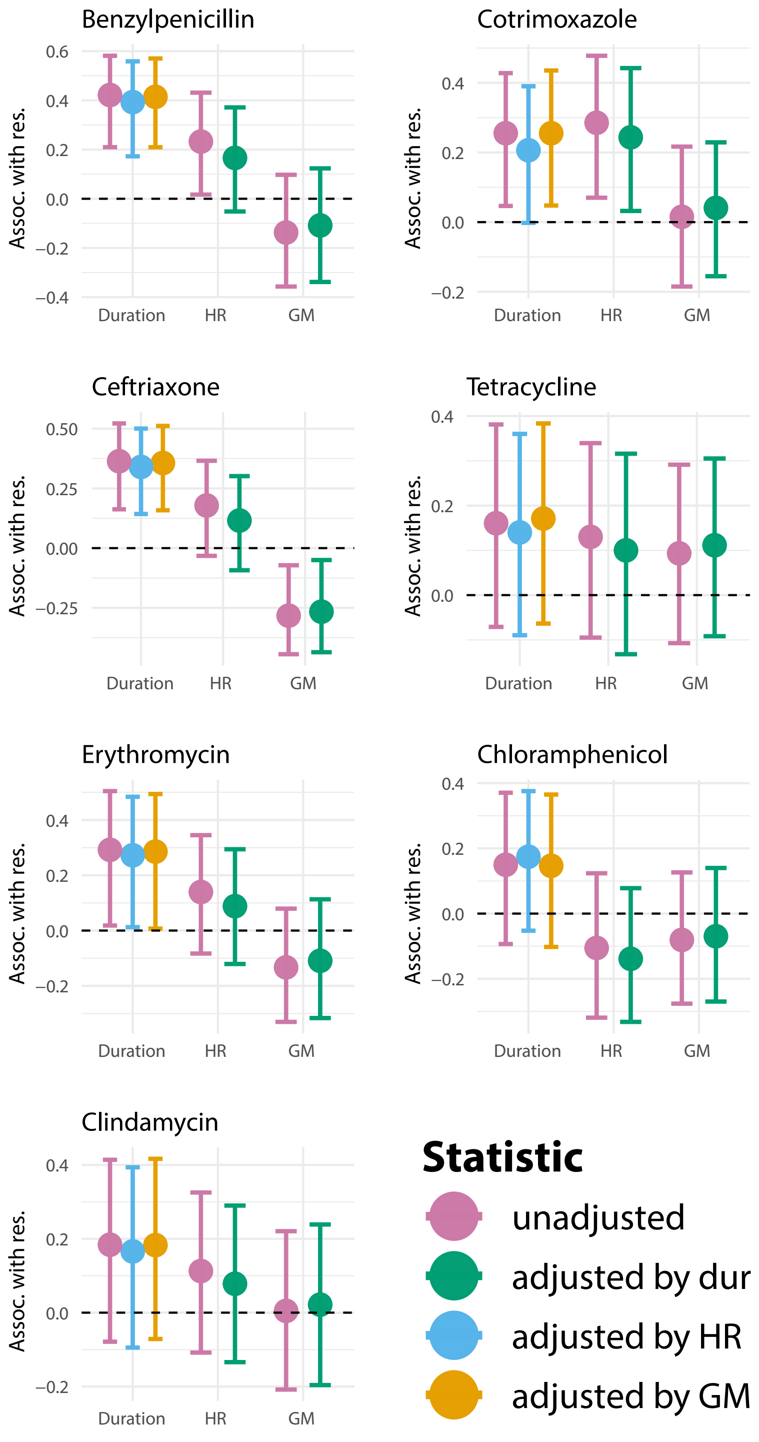


**Figure S5.** Results for individual classes of antibiotics with serotype-SC as the main unit of analysis. The legend of each panel is the same as described in Figure 2. Number of bootstrap samples is $n_b = m_b = 100$.

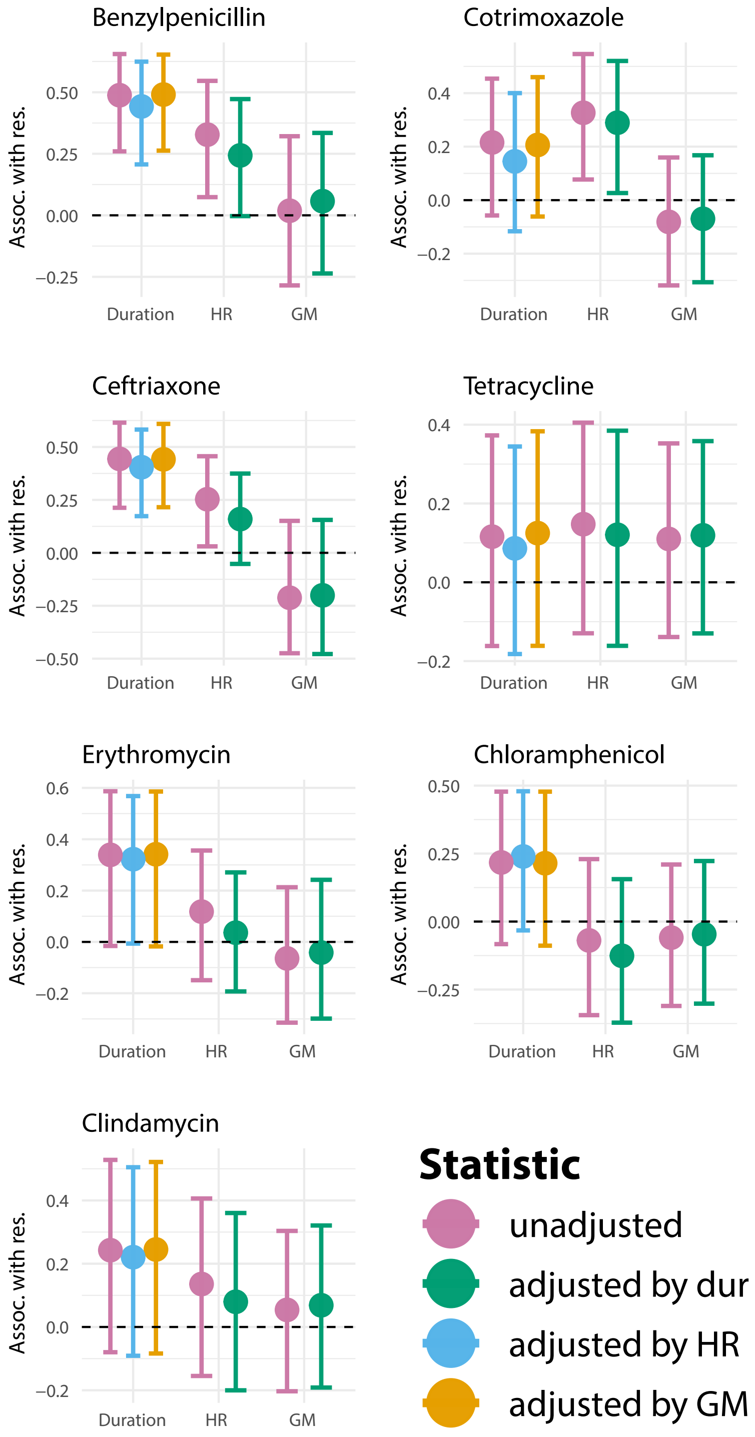


**Figure S6.** Results for individual classes of antibiotics with serotype as the main unit of analysis. The legend of each panel is the same as described in Figure 2. Number of bootstrap samples is $n_b = m_b = 100$.

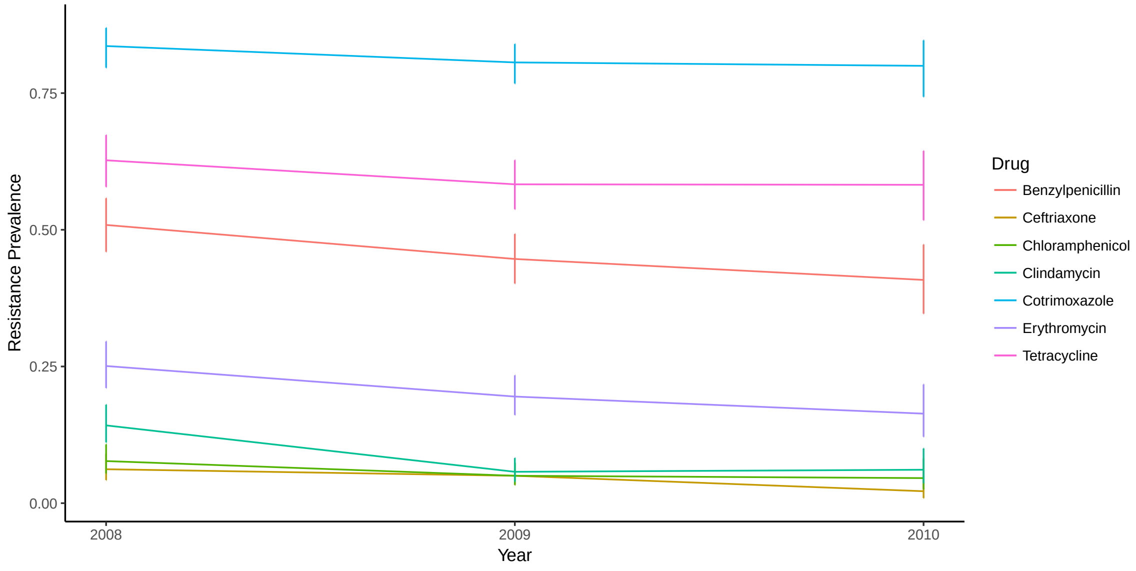


**Figure S7.** The prevalence of resistance against the seven different antibiotics as a function of time in the Maela population. Error bars represent 95% confidence intervals. For clarity, data from 2007 has not been plotted as only 6 carriage episodes were available.

A

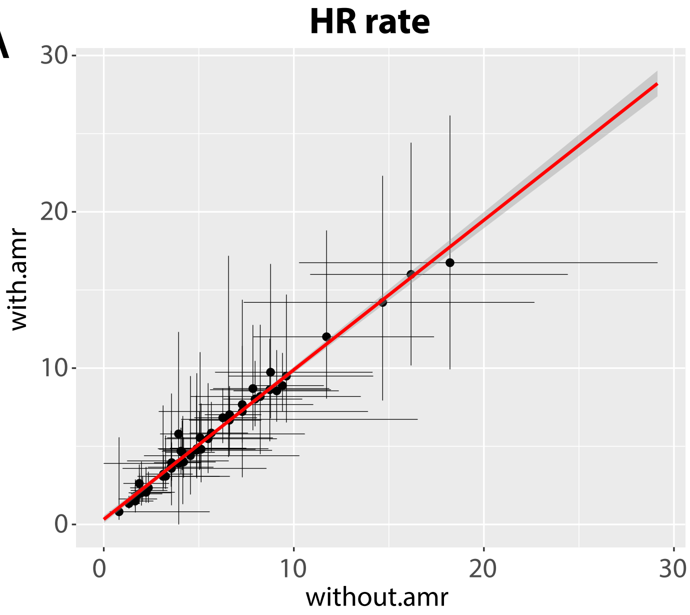


B

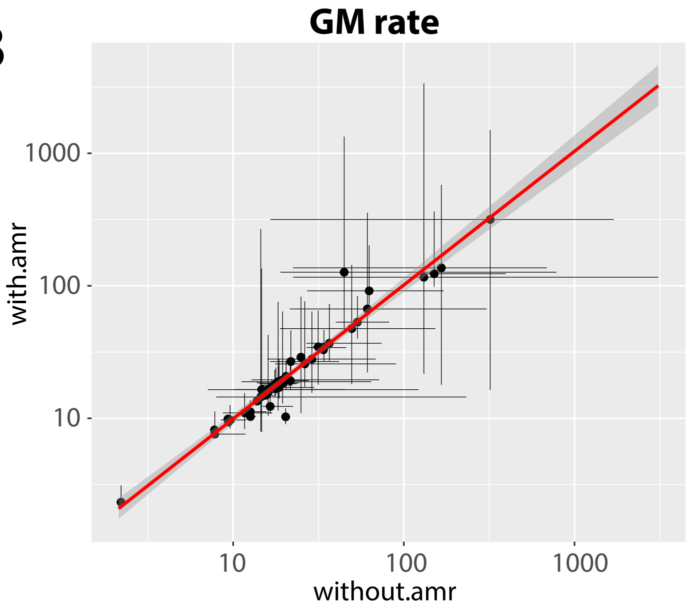


**Figure S8.** Effect of removing genetic determinants of antibiotic resistance on estimates of the HR rate (A) and GM rate (B). Y-axis shows the values estimated in the main analysis, with the 95% confidence intervals, while X-axis shows the values re-estimated having removed antibiotic resistance determinants. Such determinants were defined as COGs with representatives which gave sequence similarity (`blastp`) hits to the reference proteins defined by the ARDB database [13], using e-value of $10^{-10}$ as a cut-off.
